# Whole-brain white matter organization, intelligence, and educational attainment

**DOI:** 10.1101/297713

**Authors:** J. Bathelt, G. Scerif, K. Nobre, D.E. Astle

**Affiliations:** MRC Cognition and Brain Sciences Unit, University of Cambridge, United Kingdom; Department of Experimental Psychology, University of Oxford, United Kingdom; Oxford Centre for Human Brain Activity, Department of Psychiatry, University of Oxford, United Kingdom

**Keywords:** child development, connectome, education, intelligence, white matter

## Abstract

General cognitive ability, sometimes referred to as intelligence, is associated with educational attainment throughout childhood. Most studies that have explored the neural correlates of intelligence in childhood focus on individual brain regions. This analytical approach is designed to identify restricted sets of voxels that overlap across participants. By contrast, we explored the relationship between white matter connectome organization, intelligence, and education. In both a sample of typically-developing children (N=63) and a sample of struggling learners (N=139), the white matter connectome efficiency was strongly associated with intelligence and educational attainment. Further, intelligence mediated the relationship between connectome efficiency and educational attainment. In contrast, a canonical voxel-wise analysis failed to identify any significant relationships. The results emphasize the importance of distributed brain network properties for cognitive or educational ability in childhood. Our findings are interpreted in the context of a developmental theory, which emphasizes the interaction between different subsystems over developmental time.

## Introduction

The ability to reason and solve novel problems is a key human ability, necessary for adapting and learning from a dynamically changing environment. There is a long and rich history of philosophical and scientific exploration of the nature of human intelligence. Indeed, there are many valid ways of defining intelligence In psychology, intelligence has emerged as a core construct (Deary 2012) and understanding individual differences in intelligence has become a central focus across many subfields, including cognitive neuroscience and developmental psychology. In many cases this has a very practical application - intelligence appears to play a key role in important real-world outcomes, notably school progress (Deary et al. 2007). Intelligence correlates highly with children’s educational attainment (0.4-0.7, Mackintosh et al. 2011).

One popular way to measure intelligence is as the shared variance across multiple cognitive tasks (Spearman 1904). In this framework, intelligence is conceptualized as a multi-component system that describes individual variation in the ability to reason, solve problems, and think abstractly (Gottfredson 1997). While the relative ranking of individuals on this measure remains remarkably stable over the lifespan (Deary et al. 2000), absolute differences are amplified over the course of child and adolescent development (McArdle et al. 2002). A strong possibility is that small initial differences in one aspect of intelligence support increases in other aspects (Ferrer and McArdle 2004; Ferrer et al. 2007). Accordingly, differences in cognitive ability may contribute to better educational attainment.

Studies on the neural basis of intelligence have shifted from an emphasis on a small number of brain regions to investigations of whole-brain properties. This has in part been driven in part by new developments in analysis and conceptualization of brain structure and function. Traditional analyses have been focused on detecting local differences that result from brain insults or plasticity and were well aligned with classic localist theories that link such insults or growths/prominences to selective impairments in cognitive function (e.g. Luria, 1966; Wernicke 1874). Studies that employed voxel-wise approaches implicated the dorsolateral prefrontal cortex, anterior cingulate cortex, parietal lobe, and medial temporal cortex as the loci of intelligence (Duncan et al., 2000; Jung & Haier, 2007). In contrast, recent parallel theories in neurocognitive development and network neuroscience have emphasized the role of *interactions among* distributed brain regions (Bassett & Sporns, 2017; Colom et al., 2010; Johnson et al. 2011). According to this view, cognitive capacity emerges from the contributions of distributed brain regions that function together as an integrated network (Barbey, 2018). Recent advances that capitalize on graph theory, a branch of mathematics concerned with the study of complex systems with interacting elements, indicate that organizational principles of the whole-brain network are strongly linked to general intelligence. In network analysis, the brain is described as a set of nodes, typically brain regions, that are linked through edges that either present white matter connections or statistical associations between brain signals (Rubinov and Sporns 2010). A consistent finding is that functional and structural brain networks with higher global efficiency, i.e. networks with shorter connections between any pair of nodes in the network, are associated with higher scores on assessments of general intelligence in both children and adults (Santarnecchi et al. 2017; Pineda-Pardo et al. 2016; Kim et al. 2016; van den Heuvel et al. 2009).

Despite these recent advances in methods for exploring principles of brain organization, these are rarely applied to developmental populations. The vast majority of studies employ methods reliant upon voxel overlap and mass univariate comparisons. This can give the impression that focal differences in brain structure are associated with particular cognitive differences in childhood, and that cognitive difficulties stem from restricted lesion-like effects. However, as noted above, a strong conclusion from developmental theory is that cognitive difficulties that emerge over time are unlikely to be the result of lesion-like effects, but instead should reflect an emergent property of a dynamic system comprised of interacting subcomponents, with difficulties cascading through the system, or being partially compensated for elsewhere. It is difficult to identify these kinds of effects with traditional univariate tests.

The current study explored the relationships between whole-brain white matter network organization, general intelligence, and educational attainment in mid-childhood. We focus on two cohorts of children, one typically developing, recruited from mainstream education, and a second group of struggling learners referred by professionals in children’s specialist services. We focused on white matter because white matter maturation is an important aspect of post-natal brain development with a prolonged trajectory extending into the third decade of life (Lebel et al. 2008), and it has previously been linked to individual differences in cognition (Clayden et al. 2011). Our hypothesis is that graph theoretical measures of global brain organization will be strongly associated with children’s general cognitive ability, and that this ability will mediate links between brain organization and educational attainment. Furthermore, we predict this that these relationships will not be revealed by a more traditional neuroimaging approach, reliant on voxel overlap across children.

## Participants and Methods

### Participants

#### Attention and Cognition in Education (ACE)

This sample was collected for a study investigating the neural, cognitive, and environmental markers of risk and resilience in children. Children between 7 and 12 years attending mainstream school in the UK, with normal or corrected-to-normal vision or hearing, and no history of brain injury were recruited via local schools and through advertisement in public places (childcare and community centres, libraries). Participating families were invited to the MRC Cognition and Brain Sciences Unit (MRC CBU) for a 2-hour assessment, which included the assessments reported here, and structural MRI scanning. Participants received monetary compensation for taking part in the study. This study was approved by the Psychology Research Ethics Committee at the University of Cambridge (Reference: Pre.2015.11). Parents provided written informed consent and children verbal assent. A total of 89 children participated in the study. Twenty-six children were excluded because of low-quality MRI data (29%, see below for quality control criteria). The final sample consisted of 63 children (34 male, Age: mean=9.93, std=1.538, range=6-12).

#### Centre for Attention, Learning, and Memory (CALM)

For this study, children aged between 5 and 18 years were recruited on the basis of ongoing problems in attention, learning, language and memory, as identified by professionals working in schools or specialist children’s services in the community. Following an initial referral, the CALM staff contacted referrers to discuss the nature of the child’s problems. If difficulties in one or more area of attention, learning, language or memory were indicated by the referrer, the family were invited to the CALM clinic at the MRC CBU in Cambridge for a 3-hour assessment. This assessment included the assessments reported here. Exclusion criteria for referrals were significant or severe known problems in vision or hearing that were not corrected or having a native language other than English. Written parental consent was obtained and children provided verbal assent. Families were also invited to participate in MRI scanning on a separate visit. Participation in the MRI part of the study was optional and required separate parental consent and child assent. Contra-indications for MRI were metal implants, claustrophobia, or distress during a practice session with a realistic mock MRI scanner. This study was approved by the local NHS research ethics committee (Reference: 13/EE/0157). Of the full CALM sample, 197 children who participated in neuroimaging and had complete data on all assessments were included for the current analysis. Nine older children at the tail of the age distribution were excluded to focus on a narrower age range. A further 49 participants were excluded because of low-quality neuroimaging data (see below for criteria). The final sample for the analysis consisted of 139 children (90 male, Age: mean=9.35, std=1.683, range=5-13). The sample included a high proportion of boys as is frequently the case in sample recruited for difficulties in school (Russell et al., 2014).

### Assessments of cognition and educational attainment

#### Procedure

All children for whom we had cognitive data were tested on a one-to-one basis with a trained researcher in a dedicated child-friendly testing room at the MRC CBU. The battery included a wide range of standardized assessments of cognition and educational attainment. Regular breaks were included throughout the session. Testing was split into two sessions for children who struggled to complete the assessments in one sitting. Measures relating to cognitive performance across different domains were included in this analysis. Tasks that were based on reaction times were not included in this analysis due to their different psychometric properties compared to the included tasks that were based on performance measures.

#### Fluid reasoning

Fluid intelligence was assessed on the Reasoning task of the Wechsler Abbreviated Scale of Intelligence, 2nd edition (Wechsler 2011). Both children in the CALM and ACE sample completed this assessment.

#### Working Memory

The Digit Recall, Backward Digit Recall, Dot Matrix, and Mr X task of the Automatic Working Memory Assessment (AWMA) (Alloway et al. 2008) were administered individually. In Digit Recall, children repeat sequences of single-digit numbers presented in an audio format. In Backward Digit Recall, children repeat the sequence in backwards order. These tasks were selected to engage verbal short-term and working memory, respectively. For the Dot Matrix task, the child was shown the position of a red dot for 2 seconds in a series of four by four matrices and had to recall this position by tapping the squares on the computer screen. In the Mr X task, the child retains spatial locations whilst performing interleaved mental rotation decisions. These tasks were selected to engage visual short-term and working memory, respectively. These assessments were the same in the CALM and ACE sample.

#### Educational attainment

For the ACE sample, tasks from the non-computerized version of the Woodcock-Johnson Test of Achievement, 4th edition (WJ-IV) were administered (*Woodcock-Johnson IV Test of Achievement* 2014). ‘Letter-Word Identification’ required the reading of letters initially, with the later stages requiring full word reading. ‘Passage Comprehension’ required children to comprehend the semantic context of simple phrases initially (e.g. ‘the cat sat on the mat’, with the children shown a set of pictures), and choose the missing word for longer phrases and passages in later stages of the test. Both assessments employed “discontinue rules” to identify the child’s upper limit of ability. A final literacy test was ‘Reading Fluency’. Children were given a set of simple sentences (e.g. ‘the sky is green’) and asked to indicate whether they were true or false. Each child had 3 minutes to complete as many sentences as possible. Writing abilities were assessed using the ‘Spelling’ subtest. For this test, children were read words and contextual sentences and had to write them down in a booklet. We also included three subtests that measured mathematics ability. The first was ‘Calculation’ and simply required children to perform sums of increasing difficulty. The second measure of mathematics ability was ‘Maths Fluency’. Children were presented with relatively simple calculations in written form and asked to calculate the answers. They were given 3 minutes to do as many as possible.

For the CALM sample, spelling, reading, and maths measures were taken from the Wechsler Individual Achievement Test (WIAT, Wechsler, 2001). For the spelling assessment, children had to write words starting with simple phonetic words and progressing towards more difficult words with irregular spelling. For the spelling assessment, children were read words along with example sentences and had to write the words in an example booklet. For the reading assessments, children read words starting from short, phonetic words and progressing towards rarer, polysyllabic words. For the math assessment, children had to solve math problems starting with simple retrieval of numeric facts and progressing towards more difficult multi-stage calculations. Correct responses were scored and the assessment was terminated following the rules of the assessment manual.

**Table 1:** Scores normalized according to the mean and standard deviation of the normative sample. The statistics tested if the observed values differed from the age-expected values with a mean of 0.

| Sample | Measure | mean | std | *t* | *p* |
| --- | --- | --- | --- | --- | --- |
| CALM | Matrix Reasoning | -0.63 | 0.938 | -7.93 | <0.001 |
|  | Digit Recall | -0.49 | 1.157 | -5.04 | <0.001 |
|  | Dot Matrix | -0.49 | 0.956 | -6.02 | <0.001 |
|  | Backward Digit Recall | -0.60 | 0.792 | -8.87 | <0.001 |
|  | Mr X | -0.22 | 0.924 | -2.8 | 0.006 |
| ACE | Matrix Reasoning | 0.00 | 0.890 | 0.03 | 0.978 |
|  | Digit Recall | 0.31 | 0.795 | 3.12 | 0.003 |
|  | Dot Matrix | 0.42 | 0.936 | 3.58 | 0.001 |
|  | Backward Digit Recall | 0.29 | 0.986 | 2.34 | 0.022 |
|  | Mr X | 0.47 | 1.051 | 3.56 | 0.001 |
| CALM | Spelling | -1.17 | 0.886 | -15.57 | <0.001 |
|  | Reading | -0.96 | 1.178 | -9.64 | <0.001 |
|  | Maths | -1.05 | 1.141 | -10.89 | <0.001 |
| ACE | Letter-Word Identification | 0.66 | 0.801 | 6.52 | <0.001 |
|  | Spelling | 0.50 | 1.037 | 3.84 | <0.001 |
|  | Passage Comprehension | -0.10 | 0.780 | -0.99 | 0.326 |
|  | Reading Fluency | 0.29 | 1.065 | 2.14 | 0.036 |
|  | Calculation | 0.26 | 1.088 | 1.86 | 0.067 |
|  | Math Fluency | -0.39 | 0.853 | -3.61 | 0.001 |

### Magnetic resonance imaging

#### MRI protocol

Magnetic resonance imaging data were acquired at the MRC CBU, Cambridge U.K. All scans were obtained on the Siemens 3 T Tim Trio system (Siemens Healthcare, Erlangen, Germany), using a 32-channel quadrature head coil. For ACE, the imaging protocol consisted of two sequences: T1-weighted MRI and a diffusion-weighted sequence. For CALM, the imaging protocol included an additional resting-state functional MRI (rs-fMRI) sequence. T1-weighted volume scans were acquired using a whole brain coverage 3D Magnetisation Prepared Rapid Acquisition Gradient Echo (MP-RAGE) sequence acquired using 1mm isometric image resolution. Echo time was 2.98ms, and repetition time was 2250ms. Diffusion scans were acquired using echo-planar diffusion-weighted images with a set of 60 non-collinear directions, using a weighting factor of b=1000s*mm^-2^, interleaved with a T2-weighted (b=0) volume. Whole brain coverage was obtained with 60 contiguous axial slices and isometric image resolution of 2mm. Echo time was 90ms and repetition time was 8400ms.

#### MRI quality control

Participant movement may significantly affect the quality of MRI data and may bias statistical comparisons. Several steps were taken to ensure good MRI data quality and minimize potential biases of participant movement. First, children were instructed to lie still and were trained to do so in a realistic mock scanner prior to the actual scan. Second, all T1-weighted images and FA maps were visually inspected by a trained researcher (J.B.) to remove low-quality scans. Further, the quality of the diffusion-weighted imaging (dwi) data were assessed by calculating the displacement between subsequent volumes in the sequence. Only dwi data with between-volume displacement below 3mm was included. Further, we used residual movement as a nuisance regressor in all analyses.

### White-matter connectome construction

The white-matter connectome reconstruction followed the general procedure of estimating the most probably white matter connections for each individual, and then obtaining measures of fractional anisotropy (FA) between regions (see Bathelt et al. 2017). The details of the procedure are described in the following paragraphs (see Figure 1 for an overview).

**Figure 1:**
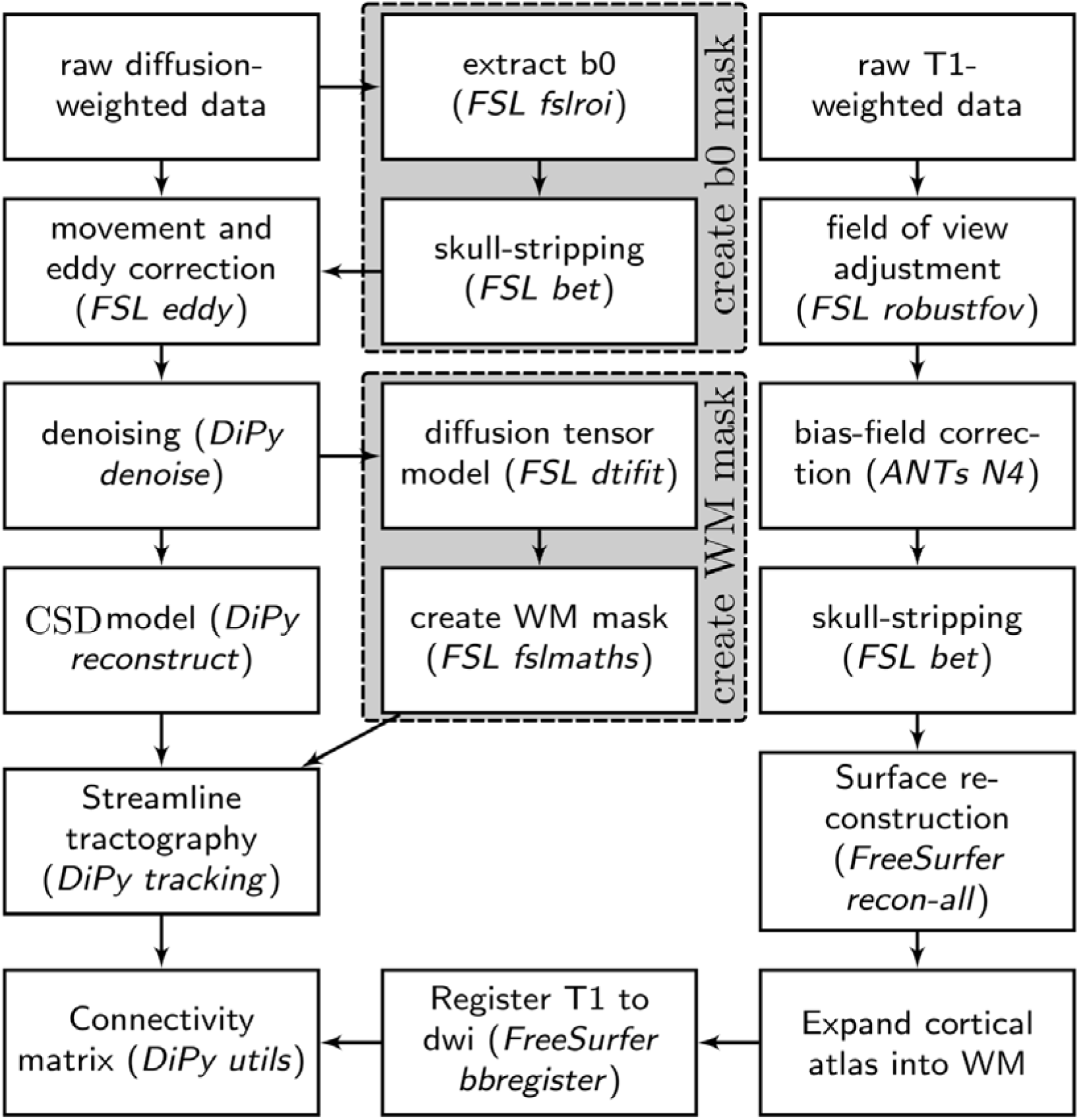
Overview of processing steps to derive the FA-weighted white matter connectome from diffusion- and T1-weighted neuroimaging data.

For the analysis, MRI scans were converted from the native DICOM to compressed NIfTI-1 format using the dcm2nii tool. Subsequently, a brain mask was derived from the b0-weighted volume of the diffusion-weighted sequence and the entire sequence was submitted for correction for participant movement and eddy current distortions through FSL’s eddy tool. Next, non-local means de-noising (Coupe 2008) was applied using the Diffusion Imaging in Python (DiPy) v0.11 package (Garyfallidis 2014) to boost the signal-to-noise ratio. The diffusion tensor model was fitted to the pre-processed images to derive maps of fractional anisotropy (FA) using *dtifit* from the FMRIB Software Library (FSL) v.5.0.6 (Behrens 2003). A spherical constrained deconvolution (CSD) model (Tournier 2008) was fitted to the 60-gradient-direction diffusion-weighted images using a maximum harmonic order of 8 using DiPy. Next, probabilistic whole-brain tractography was performed based on the CSD model with 8 seeds in any voxel with a General FA value higher than 0.1. The step size was set to 0.5 and the maximum number of crossing fibres per voxel to 2.

For ROI definition, T1-weighted images were preprocessed by adjusting the field of view using FSL’s *robustfov*, non-local means denoising in DiPy, deriving a robust brain mask using the brain extraction algorithm of the Advanced Normalization Tools (ANTs) v1.9 (Avants 2009), and submitting the images to recon-all pipeline in FreeSurfer v5.3 (http://surfer.nmr.mgh.harvard.edu). Regions of interests (ROIs) were based on the Desikan-Killiany parcellation of the MNI template (Desikan 2006) with 34 cortical ROIs per hemisphere and 17 subcortical ROIs (brain stem, and bilateral cerebellum, thalamus, caudate, putamen, pallidum, hippocampus, amygdala, nucleus accumbens). The surface parcellation of the cortex was transformed to a volume using the aparc2aseg tool in FreeSurfer. Further, the cortical parcellation was expanded by 2mm into the subcortical white matter using in-house software. In order to move the parcellation into diffusion space, a transformation based on the T1-weighted volume and the b0-weighted image of the diffusion sequence was calculated using FreeSurfer’s *bbregister* and applied to volume parcellation. To construct the connectivity matrix, the number of streamlines intersecting both ROIs was estimated and transformed into a density map for each pairwise combination of ROIs. A symmetric intersection was used, i.e. streamlines starting and ending in each ROI were averaged.

### Graph theory

The current analysis focused on local and global efficiency (E_j_, E_G_) because this metric has been found to relate closely to measures of intelligence in previous studies (Kim et al., 2016; Pineda Pardo et al., 2016). We calculated local and global efficiency for weighted undirected networks as described by Rubinov & Sporns 2010. The shortest path length between two nodes *i* and *j* in a weighted network is defined as where is a map from weight to length *and* is the shortest weighted path between *i* and *j*. The weighted global efficiency (E_G_) is defined as the average of local efficiencies (E_j_): 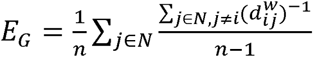 with *N* as the set of all nodes in the network and *n* the number of nodes.

Spurious connections in streamline tractography are a common problem in structural connectome studies (Zalesky et al. 2016). Typically, a threshold is applied to remove false positive streamlines. However, the choice of this cut-off is largely arbitrary. To remove the effect of setting any particular threshold, a range of density thresholds (0.3 to 0.9) was applied and the area under the curve for each graph metric was compared in subsequent analyses (van Wijk, Stam, and Daffertshofer 2010).

### Statistical analysis

Raw scores were used for the analysis after regressing the effect of other variables. For cognitive and educational attainment measures, the linear and quadratic effect of age was regressed. This allows for better control of age-related effects than using age-standardized scores calculated with reference to a normative sample, which typically apply age-standardisation with a relatively coarse resolution of several months or more. For E_G_, additional effects of brain volume as estimated through FSL SIENAX (Smith et al. 2002) and displacement during the diffusion sequence estimated through FSL eddy (Andersson and Sotiropoulos 2016) were used as additional nuisance regressors. Brain volume was included as a regressor because of the known association with intelligence (McDaniel 2005) that may create a confound for the current analysis. Displacement was included because movement during the diffusion sequence may influence measures derived from diffusion-weighted imaging (Yendiki et al. 2014). Therefore, we applied strict quality control and used volume-by-volume displacement as an additional regressor. Further, we included sex as a nuisance regressor to account for the different composition of the samples. See Table 2 for the association between the nuisance regressors and outcome variables. For detection of multivariate outliers, we calculated the Mahalanobis distance for cognitive and educational attainment variables in the ACE and CALM dataset, but no outliers were indicated at the standard cut-off. The normality assumption of regression models was checked using the Shapiro-Wilk test. A significance level of α<0.05 was used for all analyses.

**Table 2:**
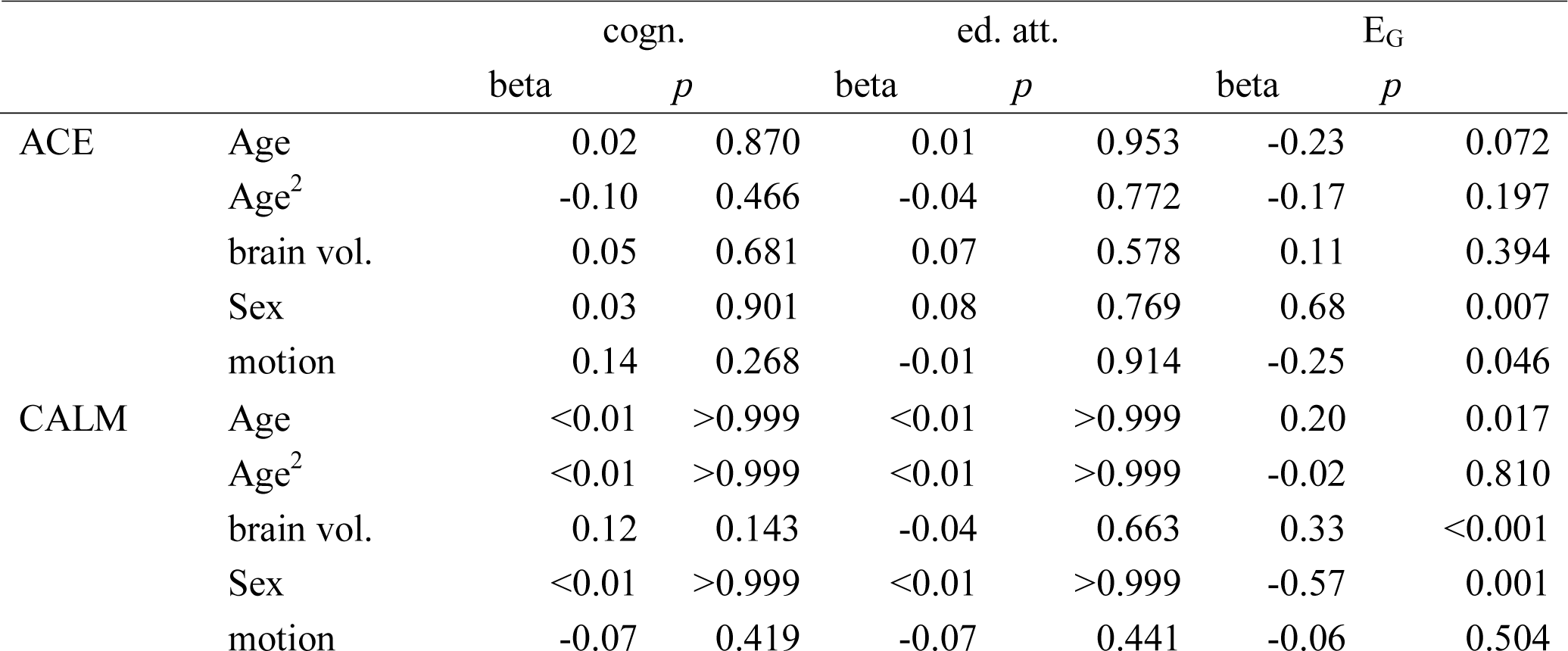
Results of simple regression between outcome variables and nuisance variables

Regression analyses were carried out using the scientific python (SciPy) v0.18.1 and *statsmodels* v0.6.1 packages for Python. Visualizations were created using the *matplotlib* v1.5.1 package for Python. The principal component analyses (PCA) were carried out using the *psych* package v1.7.2 for R 3.4.3, which uses singular-value decomposition to determine principal components. No factor rotation was applied. The mediation analyses were performed using the *lavaan* package v0.5 in R (Rosseel 2012).

### Comparison analysis with tract-based spatial statistics

To contrast the structural connectome approach with more commonly used voxelwise statistical analysis, FA maps were processed using tract-based spatial statistics (TBSS) as implemented in FSL v5.0.9. (see Smith et al. 2006 for detailed description of TBSS). In short, FA maps were moved to common space via affine and non-linear transformations using FSL tools. A common template based on a large developmental sample constructed using advanced normalization tools (ANTs) v1.9 (Avants 2011) was used as the registration target (NKI Rockland Enhanced Sample, Nooner et al., 2012). Next, the mean FA image was created and thinned to create a mean FA skeleton which represents the centers of all tracts common to the group. Each subject’s aligned FA data was then projected onto this skeleton.

For statistical comparison, the positive and negative association between FA values and cognitive or educational attainment factor scores was evaluated controlling for the effect of age, sex, movement, and intracranial volume in a generalized linear model (GLM). The model also contained an intercept term. Inflation of error rates due to multiple comparison across voxels was controlled using cluster-free threshold enhancement as implemented in FSL randomise (Winkler 2014).

## Results

### Factor analysis of cognitive and educational attainment measures

We applied principal component analysis (PCA) to investigate the factor structure of cognitive and educational attainment assessments. Established indicators were used to investigate the suitability of the age-regressed assessment scores for PCA. For both cognitive and educational attainment measures, all assessments scores correlated with 0.3 or above with at least one other assessment score for the ACE and CALM sample indicating reasonable factorability (see correlation matrices Figure 2). Further, the Kaiser-Meyer-Olkin metric indicated a sufficient measure of sampling adequacy (MSA, ACE: cognitive=0.72, educational attainment= 0.71; CALM: cognitive= 0.76, educational attainment=0.64), and Bartlett’s test of sphericity was significant (ACE: cognitive: ***χ***^2^ =71.79, df=10, *p*<0.001; educational attainment: ***χ***^2^ =219.65, df=15, *p*<0.001; CALM: cognitive: ***χ***^2^=94.37, df=10, *p*<0.001; educational attainment: ***χ***^2^=131.46, df=3, *p*<0.001). Initial eigenvalue analysis indicated that the first factor explained the highest proportion of variance for cognitive and educational attainment measures in both samples (CALM: cognitive=0.45, educational attainment=0.70, ACE: cognitive=0.55, educational attainment=0.65). Additional factors explained far less variance (see scree plots in Figure 2). Regarding factor loading, Dot Matrix and Mr X had the highest coefficients for the cognitive factor in both samples, but the loading was fairly even across assessments (see tables in Figure 2). For the educational attainment factor, all assessments loaded evenly on the educational attainment factor in both samples (see Figure 2).

**Figure 2:**
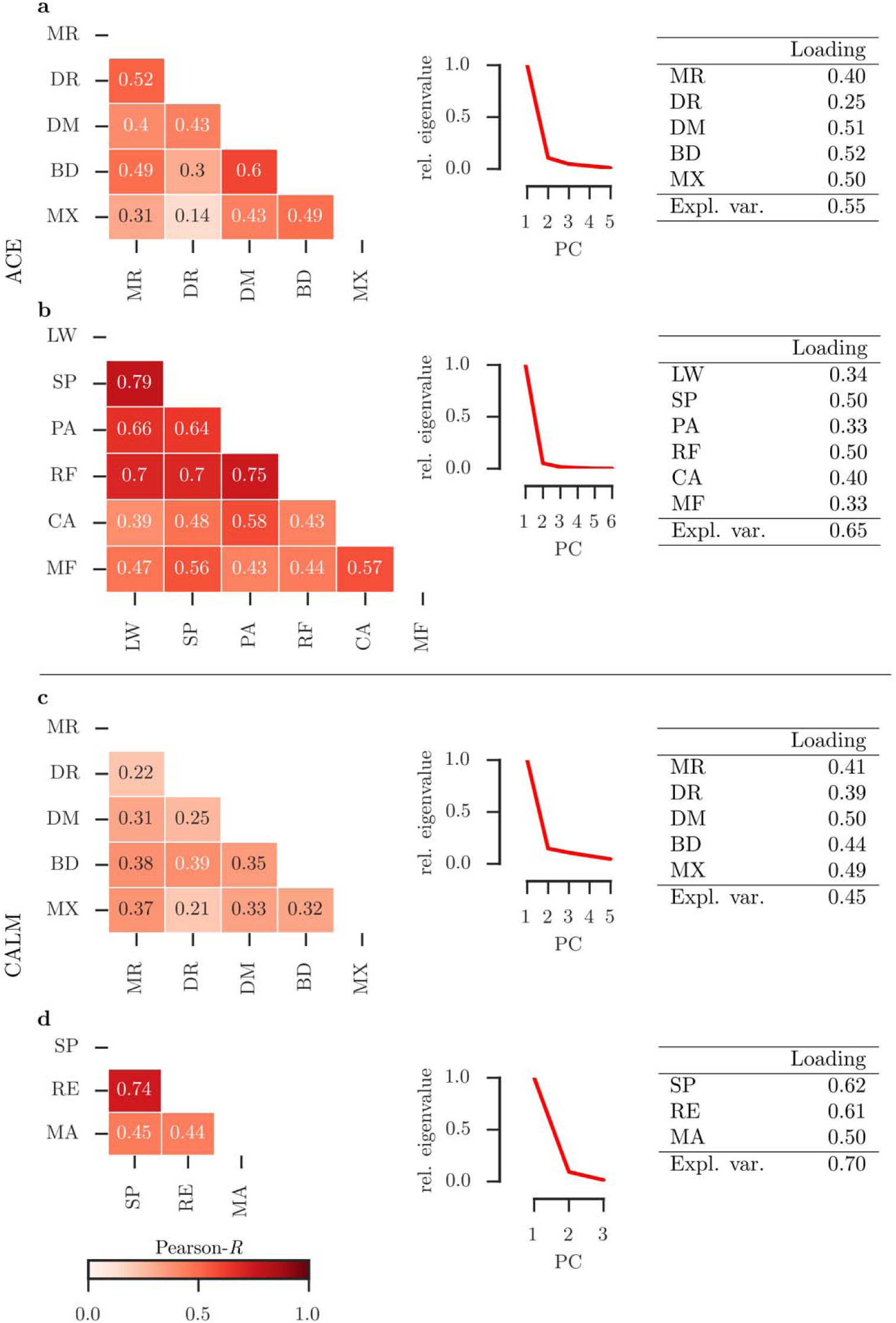
Principal component analysis (PCA) of cognitive (a,c) and educational attainment measures (b,d) in the ACE (a,b) and CALM sample (c,d). The left panel shows a matrix of correlations between measures for each factor. The middle panel shows the eigenvalues for each principal component (scree plot). The right panel shows the loading of each measure on the first principal component and the total explained variance for the first factor. Abbreviations: MR=matrix reasoning, DR=Digit Recall, DM=Dot Matrix, BD=Backward Digit Recall, MX=Mr. X, SP=Spelling, RE=Reading, MA=Maths, PC=Principal Component, Expl. var.=Explained Variance.

### Relationship between global efficiency, cognitive factor scores, and educational attainment scores

Regression analysis indicated that cognitive factor scores were a strong predictor of educational attainment factor scores (ACE: F(1,61)=25.56, R^2^=0.30, beta=0.54, p<0.001); CALM: F(1,137)=75.75, R^2^=0.36, beta=0.61, *p*<0.001) that explained between 30 and 35% of variance in educational attainment scores. E_G_ was also strongly related to cognitive factor scores in the ACE and CALM sample (ACE: F(1,61)=5.35, *p*=0.024, R^2^=0.081, beta=0.427, *p*=0.024; CALM: F(1,137)=14.32, *p*<0.001, R2=0.095, beta=0.308, *p*<0.001). Further, E_G_ was also strongly related to educational attainment scores in both sample (ACE: F(1,61)=9.048, p=0.003, R^2^=0.129, beta=0.533, *p*=0.004; CALM: F(1,137)-.16.47, *p*<0.001, R^2^=0.11, beta=0.328, *p*<0.001).

Next, we investigated the relationships between all variables in a mediation model. The results indicated that cognitive factor scores were mediating the relationship between E_G_ and educational attainment scores in both samples (see Figure 3).

**Figure 3:**
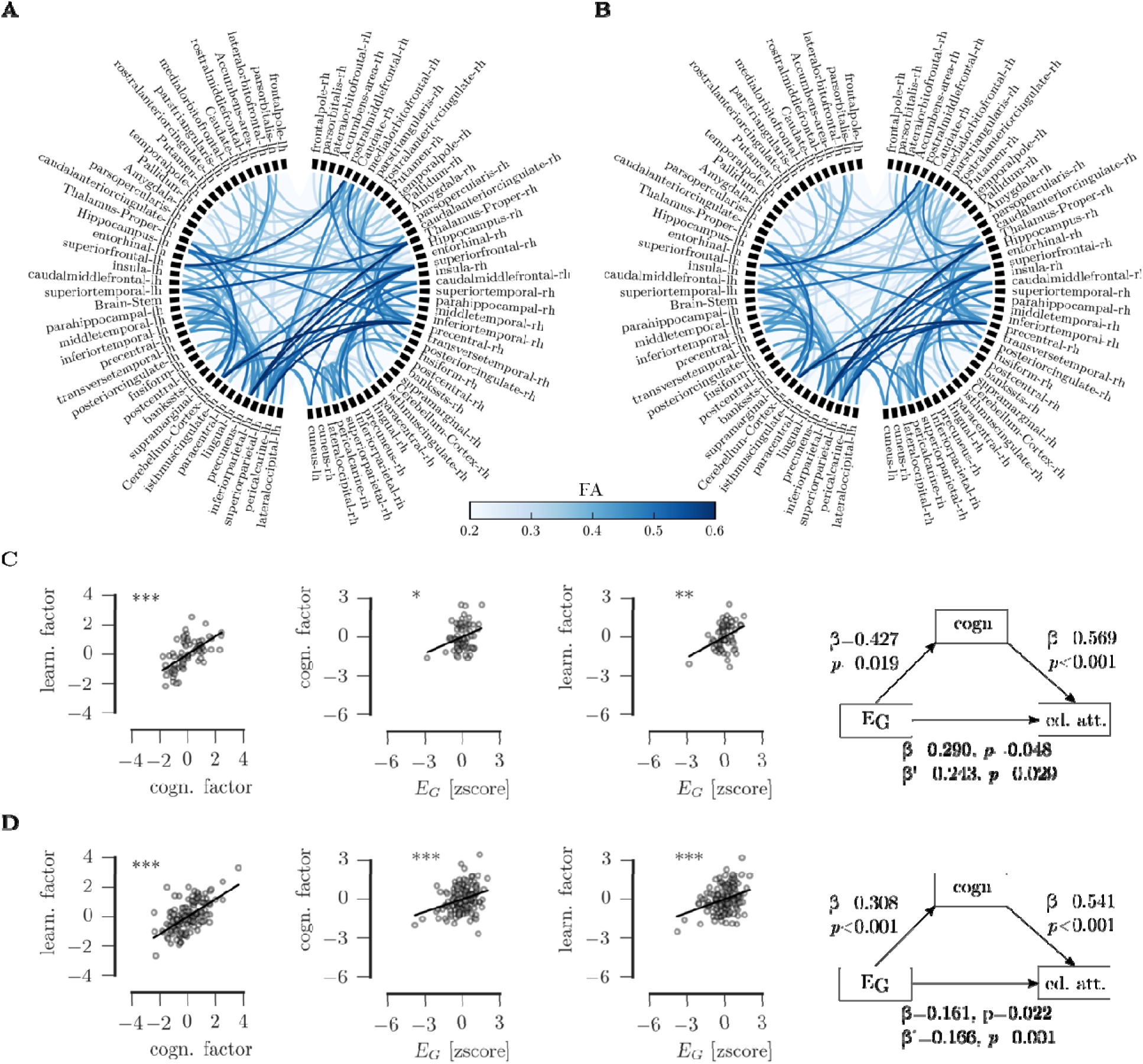
A) Visualization of the group-average connectome in the ACE sample b) Group-average connectome in the CALM sample. C) Regression of factor scores for educational attainment, cognition, and global efficiency of the white matter network and mediation relationships for the ACE sample D) Regression and mediation results in the CALM sample. Legend: β: direct effect, β’: indirect effect, ***p<0.001, **p<0.01, *p<0.05

### Regional associations

To identify brain regions that were most closely associated with cognitive and educational attainment outcomes, we assessed the strength of the linear association between local efficiency (E_j_) and factor scores in the ACE and CALM sample (see Figure 4). For cognitive factor scores, the association with E_j_ was strongest for the left anterior cingulate cortex (β=0.30), left inferior parietal cortex (β=0.27), and right rostral middle frontal cortex (β=0.29) and caudal middle frontal cortex (β=0.29) in the ACE sample, and with the left superior (β=0.31) and middle temporal cortex (β=0.28), left inferior frontal gyrus (triangular region, β=0.28), left superior parietal cortex (β=0.26), and the right superior temporal cortex (β=0.25), right lateral orbitofrontal cortex (β=0.27), and right inferior frontal gyrus (opercular region, β=0.24) in the CALM sample. For educational attainment scores, the association with E_j_ was strongest for the left inferior frontal gyrus (β=0.27), left inferior parietal cortex (β=0.28), and right rostral middle frontal gyrus (β=0.28) in the ACE sample, and the left superior (β= 0.27), middle (β= 0.26), and inferior temporal cortex (β=0.27), left inferior parietal cortex (β= 0.25), left posterior cingulate (β= 0.24), and right lateral occipital cortex (β= 0.26), righter superior parietal cortex (β=0.24), right lateral orbitofrontal cortex (β=0.23), and right temporal pole (β=0.24) in the CALM sample (see Figure 4).

**Figure 4:**
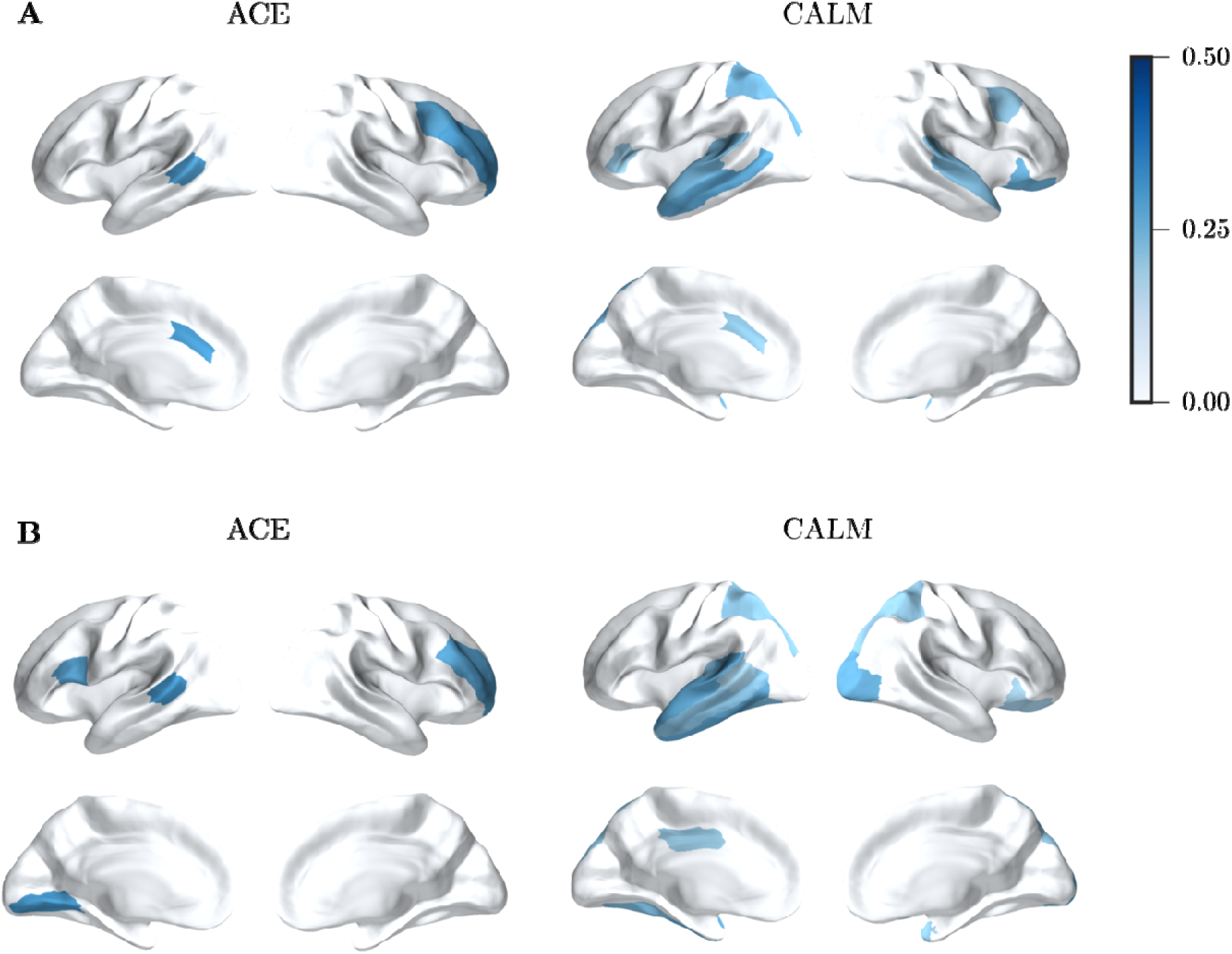
A) Strength of the association between E_J_ and cognitive factor scores in the ACE and CALM sample B) Association strength for educational attainment factor scores.

### Voxel-wise associations with cognition and educational attainment

We performed an alternative analysis to evaluate the association between voxel-wise FA and cognitive and educational attainment scores. There were no statistically significant positive or negative associations at *p*_corrected_<0.05 (see Figure 5).

**Figure 5:**
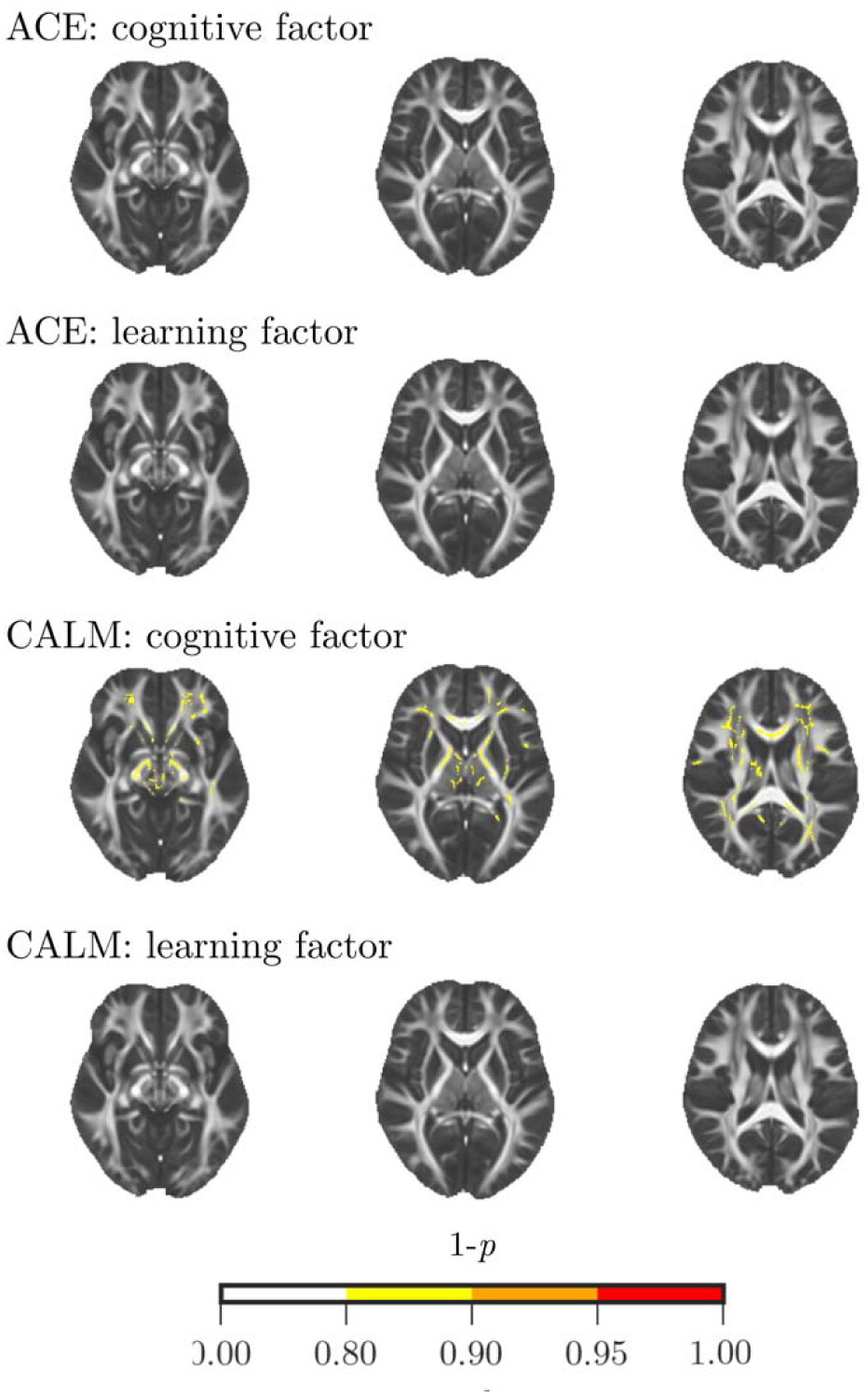
Results of voxel-wise analysis using tract-based spatial statistics (TBSS). There were no significant positive or negative associations between voxel FA values and cognitive or educational attainment measures in the ACE or CALM sample (p_corrected_>0.9). Only maps of positive associations are shown.

## Discussion

The current study investigated the relationship between cognitive ability, educational attainment, and the global and local efficiency of the white matter connectome. Global efficiency of the white matter connectome was strongly associated with children’s cognitive and educational ability - higher global efficiency was related to better fluid reasoning, both in a sample of children with age-expected ability and a sample of struggling learners.

In a first step, we derived factors for cognition and learning in our two independent samples of typically-developing children and struggling learners. For cognitive measures, a single-factor solution was favoured for both samples that explained around 50% of the variance in cognitive scores and loaded roughly equally on measures on fluid reasoning and visuospatial and verbal short-term and working memory. This factor is likely to reflect general intelligence (g) given the similarity of the current results with meta-analytic studies of this construct (Carroll 1993, Jensen 1998). However, four of the five measures used to derive the cognitive factor were short-term and working memory tasks. Intelligence and working memory are separate constructs that can be distinguished using factor analysis (Abreu 2010, Ackerman 2005), although typically using factor rotation or confirmatory factor analysis to tease the factors apart. The single-factor solution in the current study may be skewed towards a higher contribution of working memory and may only partially reflect contributions of a separate fluid intelligence construct. However, general intelligence may also be regarded as a hierarchical structure with an overarching shared variance between all cognitive measures that can be subdivided into separate domains, including short-term and working memory (Carroll 1993). This may explain the roughly equal contributions of Matrix Reasoning and the short-term and working memory tasks for the first unrotated principal component in the current analysis.

The current study also finds a high correlation between academic attainment tasks aimed at assessing reading, writing, and arithmetic that were consistently observed in two independent samples and using different standardised assessments. Factor analysis also favoured a single factor solution for the learning assessments. Reading and maths have generally been regarded as separate domains with specific mechanisms being associated with learning in each domain. This is also reflected in categorisation of learning difficulties that emphasizes specific deficits is associated with reading (dyslexia) and maths (dyscalculia). However, recent studies show a high degree of overlap that indicates that performance in one domain is highly related to performance in another (Kovacs et al., 2007). Higher correlations within domains (reading/writing vs maths) indicate that the domains may be separable, e.g. through factor rotation, but the high correlation across all tasks suggests an overriding single factor that explains a large degree of variance. The single factor for learning may reflect similarities in the assessment, i.e. standardised assessment with similar materials and instructions by the same assessor, or may reflect a common source of variance that has a similar impact on academic attainment. This common source of variance may stem from a common constraint through general intelligence as suggested by the close relationship between the cognitive and learning factors in the current analysis. In both samples, cognitive factor scores explained more than 30% of the variance in the learning factor. This finding fits in well with meta-analyses showing that fluid reasoning is closely related to school performance (Kuncel 2010, Lubinski 2009), particularly when structured assessments are used to assess academic attainment (Duckworth 2012).

In a second step, we assessed the relationship between global efficiency (E_G_) of the white matter network with general intelligence and educational attainment factor scores. The results indicated that E_G_ was related to fluid reasoning replicating findings in adults and children (Kim 2016, Haier 2004, Li 2009). This relationship was observed in a sample of children with scores within the typical range and a sample of struggling learners, which suggests that the relationship between global white matter organisation and general intelligence extends to the lower performance range. Analysis of regional associations indicated the same regions that had previously been implicated in voxel-wise analyses, i.e. the prefrontal cortex, parietal lobe, and medial temporal cortex (Jung & Haier, 2007; Deary, 2010). One possibility is that these regions play a key role in integrating whole-brain neuronal activity, acting as hubs (van den Heuvel et al. 2012). This notwithstanding, the graph analytic results suggest that properties of the whole-brain white matter connectome are more closely related to general intelligence and educational attainment than white matter integrity of any particular white matter substrate. Inferior connectivity in any part of the network may be compensated for by better connectivity elsewhere (Fornito et al., 2015), which may explain the importance of whole-brain properties and lack of overlap across individuals in the voxel-wise TBSS analysis.

It is important to bear in mind that the results of the current study come with some limitations. First, we used samples of children with typical performance and children referred for difficulties in school. It is not clear from the current analysis if associations between white matter network properties, intelligence, and educational attainment extend to superior performance. Further, both samples were cross-sectional, which precludes the analysis of age-related associations. Longitudinal data would allow us to explore how changes in structural connectomics are linked with improvements in educational attainment and cognitive ability. Another potential caveat concerns the methodology of the connectome construction in the current study. A multitude of methods for constructing structural connectomes from diffusion-weighted data have been proposed with little validation of methods through histological comparisons (Qi et al., 2015). The methods employed in the current study were chosen to reflect recommended practices (Craddock et al., 2013), but their relationship to histological measurements remains to be validated.

In conclusion, the results of the current analysis indicate that higher global efficiency of the white matter connectome is associated with better general intelligence and educational attainment in children and adolescents with performance in the age-expected and below age-expected performance. These findings support views that emphasize the importance of distributed network for higher-level cognitive processes (Johnson, 2011; Barbey, 2018).

## Acknowledgements

This work has been supported by Medical Research Council intramural programmes (MC-A0606-5PQ40 for J.H.; MC-A0606-5PQ41 to J.B. and D.A.).

The Attention and Cognition in Education (ACE) data was collected by Amy Johnson BSc as part of an MRC-funded doctoral studentship with assistance from Gemma Crickmore BSc and Erin Hawkins PhD.

The Centre for Attention Learning and Memory (CALM) research clinic is based at and supported by funding from the MRC Cognition and Brain Sciences Unit, University of Cambridge. The Principal Investigators are Joni Holmes PhD (Head of CALM), Susan Gathercole PhD (Chair of CALM Management Committee), Duncan Astle PhD, Tom Manly PhD, and Rogier Kievit PhD. Data collection is assisted by a team of researchers and PhD students at the CBSU that includes Sarah Bishop BSc, Annie Bryant BSc, Sally Butterfield MPhil MA, Erica Bottacin MSc, Lara Bridges BSc, Gemma Crickmore BSc, Fanchea Daly MSc, Laura Forde MSc, Andrew Gadie BSc, Sara Gharooni MSc, Erin Hawkins PhD, Agniezska Jaroslawska PhD, Amy Johnson PhD, Silvana Mareva MA, Sinead O’Brien MSc, Cliodhna O’Leary MSc, Joseph Rennie BSc, Ican Simpson-Kent BSc, Francesca Woolgar BSc, and Mengya Zhang MSc.

The authors wish to thank the many professionals working in children’s services in the South-East and East of England for their support, and to the children and their families for giving up their time to participate in the research.

